# Are frameworks independent from CDRs in antibodies? Exploring CDR-framework correlation networks of antibodies

**DOI:** 10.64898/2025.12.11.693675

**Authors:** Amauri Donadon Leal, Paulo Sergio Lopes de Oliveira, Leandro Oliveira Bortot

## Abstract

The growing success of monoclonal antibodies (mAbs) in therapies highlights the importance of the humanization process in antibody design. This key step involves grafting selected complementarity-determining regions (CDRs) onto human frameworks (FWs) to reduce immunogenicity. However, the effect of this process on the antibody’s structure and dynamics is underexplored. This study uses molecular dynamics simulations to evaluate whether different CDRs can impact the framework of the antibody. Pertuzumab and Trastuzumab, two clinically approved antibodies, were selected for this evaluation because they differ only by their paratope. We performed molecular dynamic simulations of the Fab portion of both antibodies in antigen-bound and unbound conditions to analyze their interdomain mobility, orientation, interchain contacts and CDR-FW dynamical correlation networks. Our results show structural effects induced by distinct CDRs. Pertuzumab displayed greater domain mobility and distinct elbow angle distributions, particularly in the heavy chain. Additionally, interchain contacts differed between the antibodies, involving both CDR and FW residues. Cross-correlation analyses further revealed distinct CDR-FW communication networks, demonstrating long-range structural impacts of CDR residues on the elbow and constant domains. These findings underscore that CDRs significantly influence the antibody dynamics through CDR-FW correlation networks. This emphasizes the importance of considering these correlation networks in antibody design to enhance efficiency of the humanization process.

## INTRODUCTION

Monoclonal antibodies (mAbs) have risen as one of the most rapidly emerging classes of biopharmaceuticals, driven by their clinical success in therapies and the billions of dollars they generate in revenue for the global markets.^1,2^ Antibodies are complex glycoproteins essential for the immune system. They are composed of a crystallizable fragment (Fc) and two identical antigen-binding fragments (Fabs), which comprise residues of the heavy and light chains. Each Fab consists of variable (VH and VL) and constant (CH and CL) domains of each chain, which are connected by flexible loops known as elbow regions. Within the variable domains, the antigen-binding site, or paratope, is formed by a total of six hypervariable loops known as complementarity-determining regions (CDRs) that are supported by scaffold-like framework regions (FWs).^3,4^ Three CDR loops are in the heavy chain (CDRH1-3) and three are in the light chain (CDRL1-3). CDRs are the core of antibodies’ variability and are responsible for antigen binding, making them the primary target of optimization strategies in antibody design,^5–8^ a rational approach that has established immunoglobulins as strong candidates for therapeutic and diagnostic applications.^9,10^

The humanization of monoclonal antibodies (mAbs) is an important step in their therapeutic design. It involves grafting CDRs from non-human sources onto human frameworks to reduce immunogenicity while retaining antigen-binding specificity and is a widely applied process to make antibodies safe for clinical use.^11–14^ Choosing the most suitable framework is an important step, as mutations or recombination of heavy and light chain frameworks have been shown to impact antigen affinity, expression rates, structural stability and Fc receptor binding.^15–19^ Moreover, the elbow angle and the VH-VL domains orientation, which is driven by structural interactions between frameworks, are related to antigen binding and antibody functionality.^20–22^ However, the inverse effect – how CDRs impact antibody’s structure – remains underexplored. The lack of this information leads to uncertainties about the conformational effects and the functionality of antibody candidates during humanization and extensive CDR optimization processes. This represents a significant gap in our understanding, particularly regarding the allosteric effects and dynamic interplay between CDRs and frameworks. So, addressing this is a critical step to improve antibody developability and optimize their therapeutic potential.

To investigate if there is an interplay between CDRs and frameworks, we selected Pertuzumab and Trastuzumab as a case study. These clinically approved therapeutic antibodies share identical frameworks and constant regions, differing only in their paratopes, and show distinct conformations in their crystal structure (see Results section), allowing us to explore how different CDRs may influence the structure of antibodies during the humanization process. Thus, here we performed molecular dynamics (MD) simulations of Pertuzumab and Trastuzumab under antigen-bound and unbound conditions to investigate how differences in paratope composition influence their dynamic correlation networks and structural behavior. MD simulations in both the presence and absence of the antigen were applied to consider possible conformational changes induced by epitope recognition.

## Methods

### RMSD in crystal structures

As mentioned earlier, the first finding that supported the use of Pertuzumab and Trastuzumab in this study was the difference in their domain orientation. First, the Root Mean Square Deviation (RMSD) of both antibodies were calculated in PyMOL^23^ with a direct alignment of entire Fab portion, excluding antigens. To assess if the observed RMSD value was caused by internal modifications in domains or not, we selected their constant or variable domains separately and performed the alignment.

### MD simulation protocol

We performed molecular dynamics simulations of four systems: Pertuzumab (PDB ID: 1S78, chains C and D) and Trastuzumab (PDB ID: 1N8Z, chains A and B), each in antibody-bound and unbound conditions. MD simulations were performed using the GROMACS ^24^ package with the ff03ws force field,^25^ which is a modified version of the AMBER ff03* force field that is optimized to better reproduce protein flexibility and the affinity of protein complexes. RMSD, RMSF, angle and SASA analyses were also performed using GROMACS. The simulation box was defined using the dodecahedral geometry, with a minimum distance of 1 nm between the protein atoms and each box edge. To neutralize system charges, sodium and chloride ions were added to achieve a total concentration of ∼150 mM, and neutral pH were modeled through the appropriate protonation states of titratable residues. For system minimization, we used the Steepest Descent algorithm.^26^ System equilibration was carried out for 1 ns under the temperature of 309K and pressure of 1 bar. The production runs were carried out for 200 ns under NPT conditions (isothermal-isobaric ensemble), which is most compatible with biological systems. The Hydrogen Mass Repartitioning (HMR) method ^27^ was used to eliminate the fastest degrees of freedom in order to enable simulations with a 4 fs timestep. All trajectory analyses considered only the last 100 ns of each simulation replicate to account for equilibration time. Five independent replicates were simulated for all systems.

### Domains’ mobility analyses

For each domain mobility analysis, RMSD calculations were performed individually for the light and heavy chains in each system. Within each chain, two separate structural alignments were considered: one using the constant domain (CL for the light chain or CH for the heavy chain) and another using the variable domain (VL for the light chain or VH for the heavy chain). For each calculation, the reference structure that deviations were calculated from was the average structure of the last 100 ns of each replicate for improved accuracy.

For each alignment, two RMSD analyses were performed. The first was calculated on the aligned domain itself (intradomain RMSD) to assess the structural integrity of the domain. The second was calculated on the other domain within the same chain (interdomain RMSD) to measure the movement of one domain relative to the other. This process was applied to each of the four domains in the Fab region.

### Angles analyses

To explore the direction of interdomain movements, we analyzed four angles in each system: two dihedral angles and two hinge or elbow angles (Figure 3). The first dihedral angle, named here as VL-CH, is defined by the center of mass (COM) of VL domain, light chain elbow, heavy chain elbow and CH domain. The second dihedral, named VH-CL, is the mirrored version of the first one, starting with the COM of VH and ending with the COM of CL. The COM of the elbow was defined as the COM of the last residue of the variable portion and the first residue of the constant portion. The segmentation of the antibody between variable and constant portions was defined according to IMGT numbering scheme ^28^ with ANARCI program.^29^ These dihedrals were inspired by previous description ^20^ and were explored to analyze the pendular movement of the variable domains.

The elbow angle of light (EL) and heavy (EH) chains were also measured to investigate the “contraction” of Fab portion. These angles are defined by the COM of the variable domain, COM of the elbow and COM of the constant domain for each chain.

As a comparison, we used ABangle ^30^ to evaluate variable domains orientations. It calculates six metrics, three of which capture similar movements to those angles we have defined above.

Statistical analyses were conducted to compare antibodies and angles. The Kolmogorov-Smirnov (KS) test was applied to compare the data variability profiles due to the presence of multimodal distributions. Also, the Spearman correlation was employed to identify any dependencies in angular motion patterns.

### Inter-chains contacts

As we are evaluating the dynamics of the antibody domains, we investigated contacts between residues of light and heavy chains. For this, we first conducted Molecular Mechanics/Poisson-Boltzmann Surface Area (MM/PBSA)^31^ calculations to identify amino acids with the highest contributions to Fab chain interactions. MM/PBSA is a widely used approach due to its computational efficiency in estimating relative binding free energies in protein-protein interactions and performing the decomposition analysis of free energy terms at the level of individual residues^32,33^. The amino acids that contributed with ΔG < −2.5 kcal/mol were selected for contact mapping. Each identified residue was annotated according to its antibody region (CDR, framework or constant portions) based on the IMGT numbering scheme^28^ and determined with AbNumber package (https://github.com/prihoda/AbNumber), which is based on the ANARCI algorithm.^29^

Subsequently, we investigated the contact partners for each identified residue. All amino acids in the opposite chain within a 3 Å distance cutoff (considering all atoms) were selected. The contact partners were also annotated according to their antibody region and the frequency at which they satisfied the distance cutoff throughout the MD simulation. This inter-chain interaction analysis was conducted using a custom-developed python script.

### Cross-correlation analyses

To evaluate the impact of CDRs over the full Fab, we calculated the Dynamical Cross-Correlation Matrix (DCCM)^34^ using the R package Bio3d,^35^ which calculates the correlation coefficient (C) between the displacements of atoms *i* and *j* throughout simulation frames in Cartesian coordinates. It results in a *n* x *n* matrix where *n* is the number of residues of the analyzed protein with C_ij_ values spanning from −1 (anticorrelated residues) to 1 (correlated residues), with 0 meaning no correlation. The correlations with the highest intensities (C_ij_ > |0.8|) that involves CDR residues were traced in the structure with PyMOL^23^ for visualization using a custom-developed python script. For this analysis, we sampled the trajectories every 200 ps.

## Results

### Antibodies have different interdomains mobility

Despite differing only in the paratope residues, the crystallographic structures of Pertuzumab and of Trastuzumab show that their domains are oriented differently (Figure 1). The overall Cα RMSD between the two structures upon superposing the backbone of the entire Fab is 3.303 Å. In contrast, the Cα RMSD is reduced to 0.891 Å and 0.902 Å when superposing the backbone of their constant and variable regions separately, indicating that the primary difference is caused by domain orientation rather than internal structural changes.

**Figure 1:**
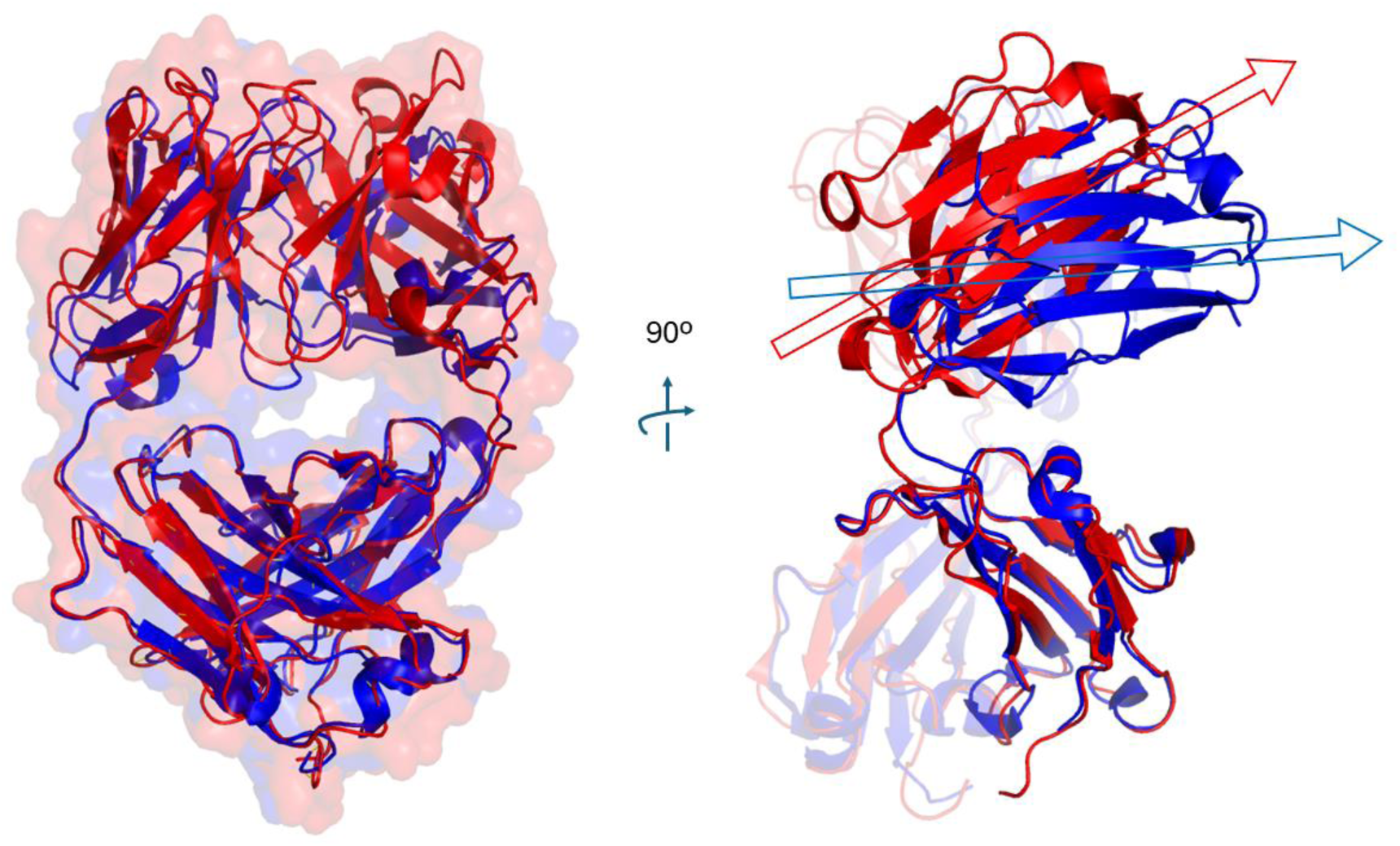
Difference in the orientation of variable domains of Pertuzumab (red) and Trastuzumab (blue) when the backbone of their constant domains is superposed. PDB IDs used: 1S78 and 1N8Z, respectively.

To further investigate such differences, we performed MD simulations to assess the domain mobility of both antibodies. Figure 2 illustrates the RMSD profiles from simulations of Pertuzumab and Trastuzumab. We observe that the intradomain RMSD values of both antibodies remain predominantly below 2 Å, indicating that each domain maintains structural stability and does not unfold during the simulations. In contrast, the interdomain RMSDs show higher median values (Supplementary Table S1) with broader distributions of Pertuzumab compared to Trastuzumab. This reflects the flexibility of these domains relative to one another and suggests that the structural dynamics of these antibodies, that differ only by their paratope, are different.

**Figure 2.**
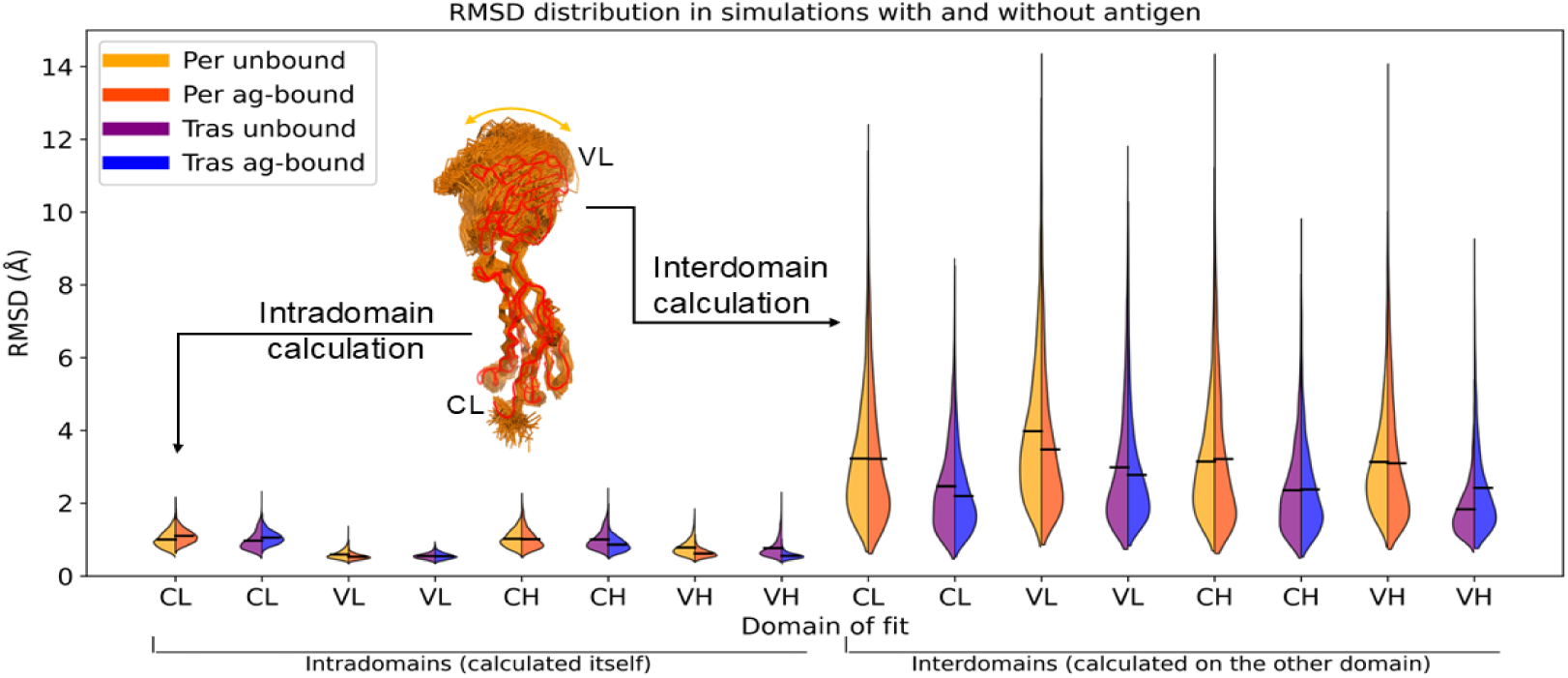
RMSD values calculated from MD simulations of Pertuzumab and Trastuzumab in antigen-bound and unbound conditions. Simulations are named with the prefix of the antibody (“per” or “tras”) in the presence or not of the antigen. The RMSD values of intradomain and interdomain calculations are shown as violin plots with an illustrative representation of the calculation of these movements.

### Differences between antibodies are due to CDR influence

Many factors may contribute to the discrepancy observed above, including (i) crystal packing, (ii) antigen-binding effects, and (iii) our hypothesis that CDRs influence internal correlation networks. To address these possibilities and clarify the direction of movement reflected in the RMSD data, we analyzed four angles related to the domain’s orientation throughout the MD simulations (Figure 3).

**Figure 3.**
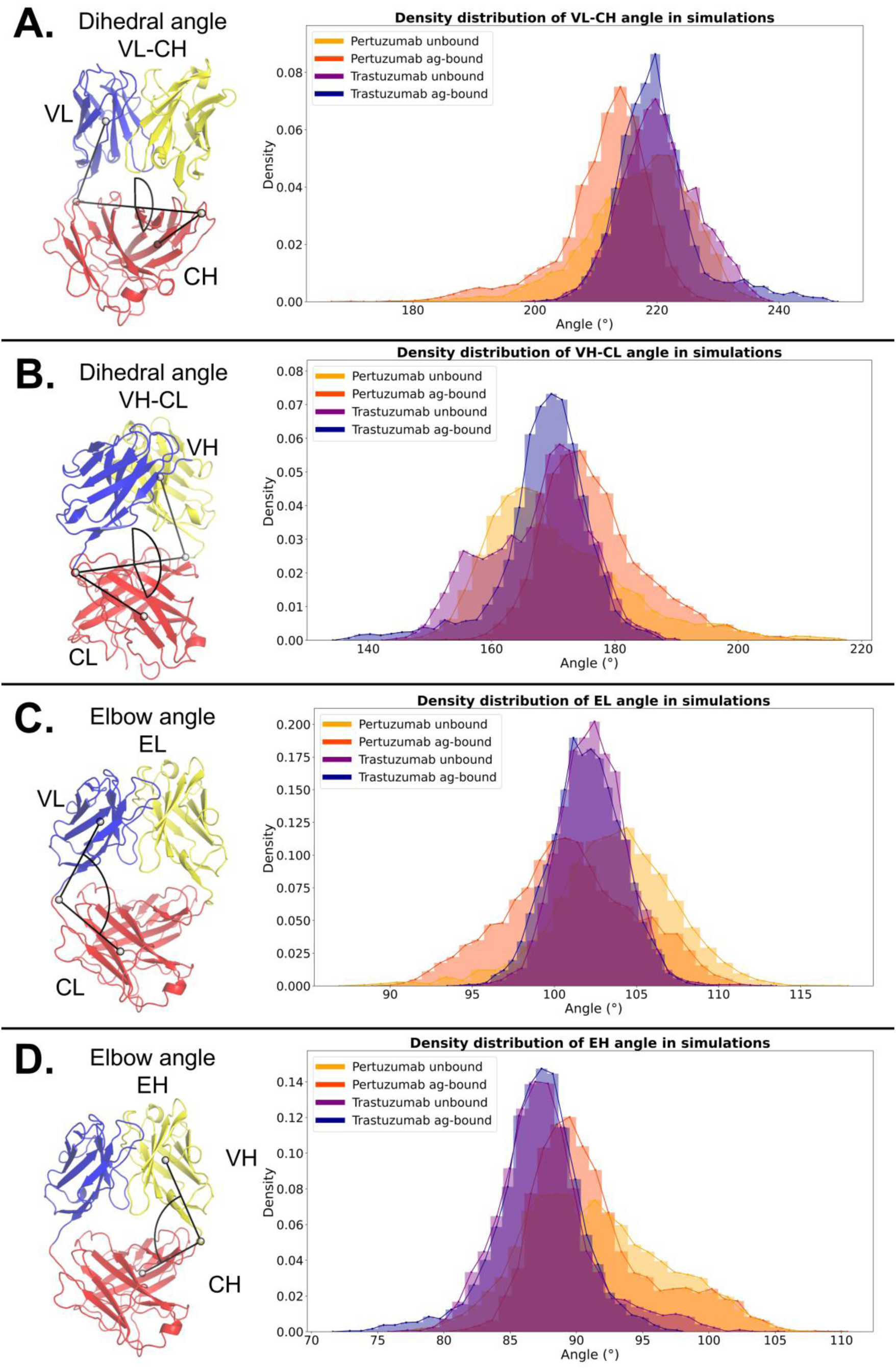
Angle profiles sampled during MD simulatios of Pertuzumab and Trastuzumab under antigen-bound and unbound conditions. Illustrative representations of VL-CH (A), VH-CL (B), EL (C) and EH (D) are shown to the left side and the density distributions sampled for each angle to the right side. Simulations are named with the prefix of the antibody (“per” or “tras”) in the presence or not of the antigen.

If crystal packing were responsible for the observed structural differences, MD simulations, which are free from such constraints, should have sampled similar angle distributions. Instead, we observed distinct angle profiles for Pertuzumab and Trastuzumab under the antigen-bound conditions, with statistically significant differences (Supplementary Table S2). The most evident differences were observed in elbow angle distributions. Notably, the EH angle (see Methods section) consistently showed the greatest divergence between antibodies in both the antigen-bound and unbound conditions. The persistence of these specific angular signatures throughout the simulations discards crystal packing as the cause of domain-orientation differences.

If antigen binding were responsible for the differences in the crystallographic structures, MD simulations should have sampled similar angle distributions for both antibodies in the unbound state. Despite some overlap in the sampling of dihedral angles, the distribution profiles remained different between the antibodies. Also, the elbow angles showed consistent statistical differences in both bound and unbound conditions (Supplementary Table S2), with emphasis on the EH angle (KS_unbound_ = 0.37, p < 0.0001; KS_ag-bound_ = 0.39, p < 0.0001), suggesting it is a key factor to describe the distinct flexibility profiles of the domains. It indicates that the antigen binding is not likely to be the main cause for the difference in domain orientation observed in the crystallographic structures. Interestingly, these results also reveal differences in the sensitivity of antibodies to antigen-induced structural changes.

A correlation analysis was performed across all measured angles. Notably, the VL-CH and VH-CL angles displayed strong negative correlations (Supplementary Tables S3 and S4), indicating compensatory antiparallel movements between the variable domains in both antibodies. Also, the EH angle was strongly correlated with both dihedral angles, highlighting its central role in antibody dynamics. In contrast, the remaining angle combinations exhibited either weak or moderate correlations, suggesting a complex landscape of independent motions of the domains. This complexity underscores the advantage of using multiple metrics to capture the diverse dynamics of each antibody, as different angles highlight unique aspects of their structural flexibility.

In addition to the method used above, the angles of variable domains were calculated by ABangle.^30^ However, the results of this method did not show statistical difference between antibodies as observed for the angles we evaluated (Supplementary Tables S5). As our angles account for constant domains, they may be more sensitive to interdomain variations, capturing information from the entire Fab. No strong correlation between the angles proposed here and those from ABangle was observed (Supplementary Tables S3 and S4), indicating that the method involving constant domain captures specific information not seen when considering only variable domains. Thus, it is reasonable to propose the angle analysis presented here as a complementary metric to ABangle, providing additional insights into the antibodies’ dynamics.

Taken together, these results indicate that neither crystal packing effects nor antigen binding induce structural changes that are large enough to override the antibody’s intrinsic conformational characteristics. Thus, it supports the hypothesis that the difference in the composition of the paratope of Pertuzumab and Trastuzumab induces long-range effects over the Fab domain.

### Inter-chains contacts

Given the findings above, we investigated the origin of the dissimilar angle distributions. To address this, we performed MM/PBSA calculations to identify residues that contribute the most to interactions between the light and heavy chains. Table 1 shows the residues that contributed with ΔG < −2.50 kcal/mol and their contact frequencies with CDR and FW residues.

**Table 1.** Residues relevant for inter-chain interactions.

| Pertuzumab |  |  |  |  |  |  |  |
| --- | --- | --- | --- | --- | --- | --- | --- |
|  |  | Unbound |  |  | Ag-bound |  |  |
| Residue<sup>a</sup> | Location<sup>b</sup> | $\Delta G$<br>(kcal/mol)<sup>c</sup> | % of contacts | | $\Delta G$<br>(kcal/mol)<sup>c</sup> | % of contacts | |
|  |  |  | with CDR | with FW |  | with CDR | with FW |
| TYR 91 | CDR3 (L) | -3.09 | 99 | 0 | > -2.5 | - | - |
| PHE 98 | FW4 (L) | -4.05 | 78 | 100 | -4.05 | 73 | 100 |
| PHE 116 | Constant (L) | > -2.5 | NA | NA | NA | NA | NA |
| PHE 118 | Constant (L) | -4.4 | NA | NA | NA | NA | NA |
| GLN 39 | FW2 (H) | -2.86 | 0 | 99 | > -2.5 | - | - |
| LEU 45 | FW2 (H) | -3.40 | 0 | 100 | -3.28 | 0 | 100 |
| TRP 47 | FW2 (H) | -2.84 | 91 | 64 | -3.43 | 100 | 81 |
| PHE 99A | CDR3 (H) | -2.56 | 99 | 30 | > 2.5 | - | - |
| TYR 99B | CDR3 (H) | -3.87 | 75 | 100 | -2.92 | 75 | 75 |
| PHE 128 | Constant (H) | > -2.5 | NA | NA | -2.51 | NA | NA |
| LEU 130 | Constant (H) | -2.66 | NA | NA | -2.56 | NA | NA |
| PHE 172 | Constant (H) | -4.91 | NA | NA | -5.21 | NA | NA |
| PRO 173 | Constant (H) | -3.08 | NA | NA | > -2.5 | NA | NA |

| Trastuzumab |  |  |  |  |  |  |  |
| --- | --- | --- | --- | --- | --- | --- | --- |
|  |  | Unbound |  |  | Ag-bound |  |  |
| Residue | Location | $\Delta G$<br>(kcal/mol) | % of contacts | | $\Delta G$<br>(kcal/mol) | % of contacts | |
|  |  |  | with CDR | with FW |  | with CDR | with FW |
| GLN 38 | FW2 (L) | -2.54 | 0 | 99 | -2.98 | 0 | 100 |
| PHE 98 | FW4 (L) | -3.64 | 73 | 100 | -3.61 | 54 | 100 |
| PHE 116 | Constant (L) | -2.73 | NA | NA | -2.66 | NA | NA |
| PHE 118 | Constant (L) | -4.96 | NA | NA | -4.67 | NA | NA |
| LEU 45 | FW2 (H) | -2.97 | 0 | 100 | -3.05 | 0 | 100 |
| TRP 47 | FW2 (H) | -3.22 | 100 | 58 | -3.17 | 100 | 72 |
| TRP 110 | FW4 (H) | -4.03 | 0 | 100 | -3.28 | 0 | 100 |
| PHE 129 | Constant (H) | -2.91 | NA | NA | -2.77 | NA | NA |
| LEU 131 | Constant (H) | -2.61 | NA | NA | -2.66 | NA | NA |
| PHE 173 | Constant (H) | -5.03 | NA | NA | -5.09 | NA | NA |
| PRO 174 | Constant (H) | -2.51 | NA | NA | > -2.5 | NA | NA |

<sup>a</sup>: Residues numbers are in accordance with the PDB file 1S78 and 1N8Z for Pertuzumab and Trastuzumab, respectively; <sup>b</sup>: The parenthesis indicates if residue belongs to the light chain (L) or heavy chain (H); <sup>c</sup>: The standard error of the mean for each $\Delta G$ value ranged from 0.03 to 0.08. NA: Non-applicable

In Pertuzumab’s unbound simulations, the residues with the strongest contributions in the variable domains were TYR 91 (CDRL3) and PHE 98 (FW4) of the light chain, as well as GLN 39, LEU 45 and TRP 47 from FW2, and PHE 99A and TYR 99B from CDRH3 of the heavy chain. Upon antigen binding, TYR 99B (CDRH3) is the only residue from the CDR that still contributed with inter-chain interactions. These findings reveal that CDR residues also play a significant role in Pertuzumab’s inter-chain interactions, underscoring the importance of carefully considering the residues of CDR during antibody design, mainly when inter-chain interactions involve the paratope.

Trastuzumab featured GLN 38 (FW2) and PHE 98 (FW4) of the light chain and LEU 45 (FW2), TRP 47 (FW2) and TRP 110 (FW4) of the heavy chain as major contributors to inter-chain interactions of variable domains in the unbound condition. All the five residues still contributed to chain contacts in Trastuzumab’s antigen-bound simulations. Interestingly, the FW residues GLN 38 and TRP 110 are also present in Pertuzumab but were not significant for its interchain interactions, which suggests that the structural impact of changing CDRs may also influence FW conserved residues and their contribution to inter-chain interactions.

To further this investigation, we developed a script to locate the residues that are in contact with those selected by MM/PBSA calculations. We observed that FW-FW and FW-CDR interactions compose inter-chain contacts in both antibodies, while CDR-CDR contacts occur only in Pertuzumab (detailed contact information is described in Supplementary Tables S6 – S9). Notably, across all simulations, at least one CDR3 residue from one chain consistently interacted with FW residues during at least 75% of the simulation time. This indicates a direct role of CDRs in inter-chain interactions, further supporting the existence and importance of CDR-FW communication networks for antibodies’ dynamics. In addition, residues involved exclusively in FW-FW contacts were found only in Trastuzumab. This result indicates that different CDRs can also impact FW-FW communication, probably from the influence of the CDR-FW network, further highlighting the interplay between these structural regions in antibodies.

Inter-chain contacts reveal another difference between the antibodies. For Trastuzumab, binding to the antigen did not affect interchain interactions established by residues from the constant domain. On the other hand, residues PHE 116 (light chain) and PHE 128 (heavy chain) of Pertuzumab showed reduced interaction energy upon antigen binding. These observations support the hypothesis that specific information networks connect the CDRs to FW regions, which also influences constant domains and explains the differences between the mobility profile of Pertuzumab and Trastuzumab. Thus, we found that the paratope residues can directly interact with the opposite chain through CDR-CDR or CDR-FW interactions, and indirectly define FW-FW contacts. This observation supports the existence of a network of interactions which influences the antibody’s structural dynamics.

### The antibodies contain specific CDR-FW networks

With the resuinteractions anderved compelling evidence supporting the existence of intrinsic networks within the antibody domains that connect CDRs to their respective frameworks. So, we performed cross-correlation analyses to further characterize these networks. The dynamical cross-correlation map (DCCM) plots with values > |0.8| of all simulations are shown in Figure 4.

**Figure 4.**
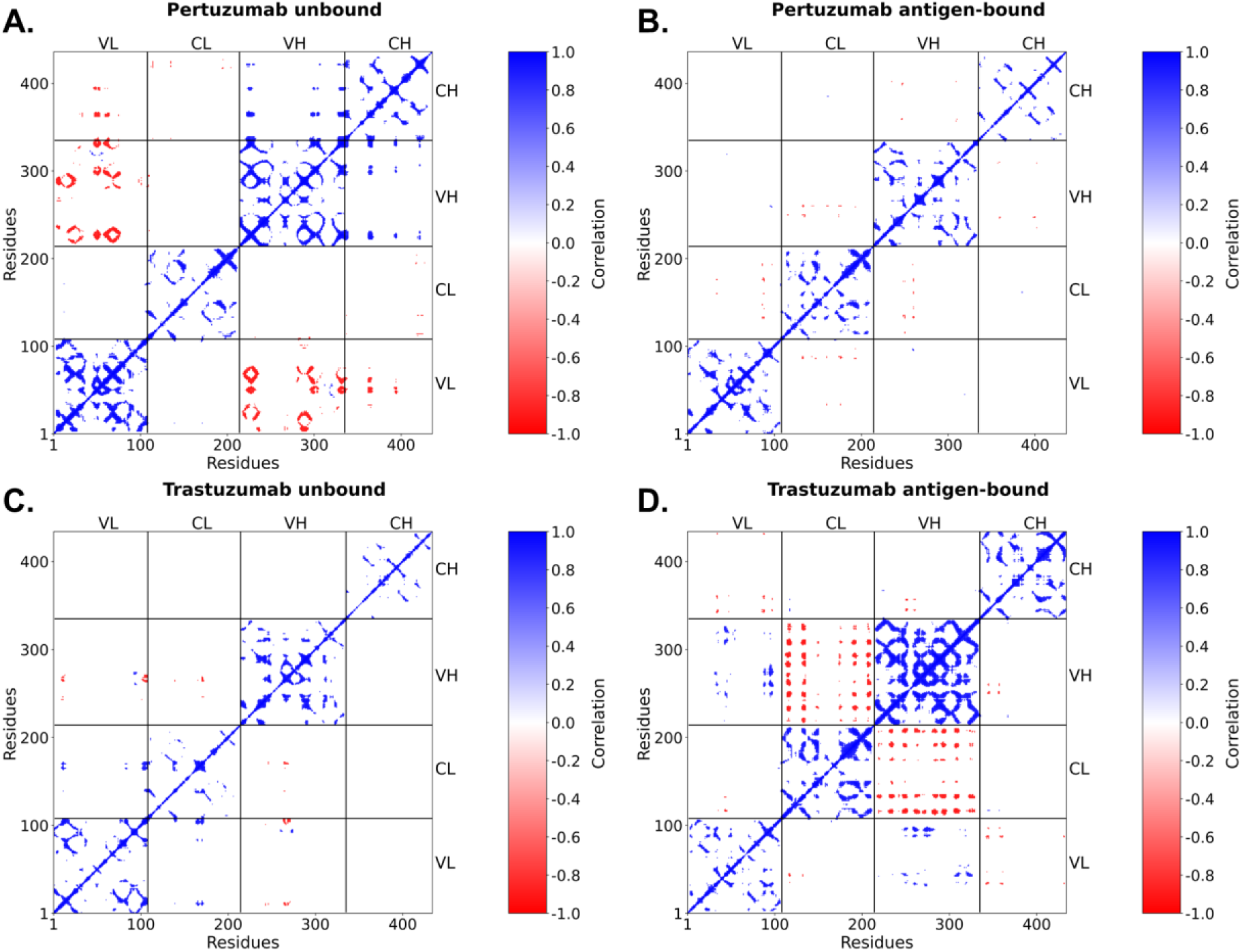
Dynamic Cross-Correlation Matrix (DCCM) plots. It shows correlations with intensity over 0.8 for Pertuzumab under unbound (A) and antigen-bound (B) simulations and for Trastuzumab under unbound (C) and bound (D) simulations. Red represents negative correlations and blue represents positive ones.

We observed that the antibodies in both unbound and antigen-bound simulations display long-range correlations across the entire Fab. Interestingly, we found differences in the strong cross-correlation pattern (C_ij_ > |0.8|) between Pertuzumab and Trastuzumab. For Pertuzumab, we observed strong long-range negative cross-correlations involving the variable domains in the unbound condition that vanished upon antigen binding. For Trastuzumab, the opposite behavior was verified.

Upon inspecting the residues that are highly correlated with CDRs, we found that the paratope residues of both antibodies have strong short-range positive correlations within the variable domain both in the antigen bound and unbound conditions (Figure 5). On the other hand, paratope residues display antibody-specific strong long-range negative correlations with residues from the constant domains. Additionally, the negative correlation network is affected by antigen binding. These results show that differences in the paratope composition can influence the dynamics of the entire Fab portion.

**Figure 5.**
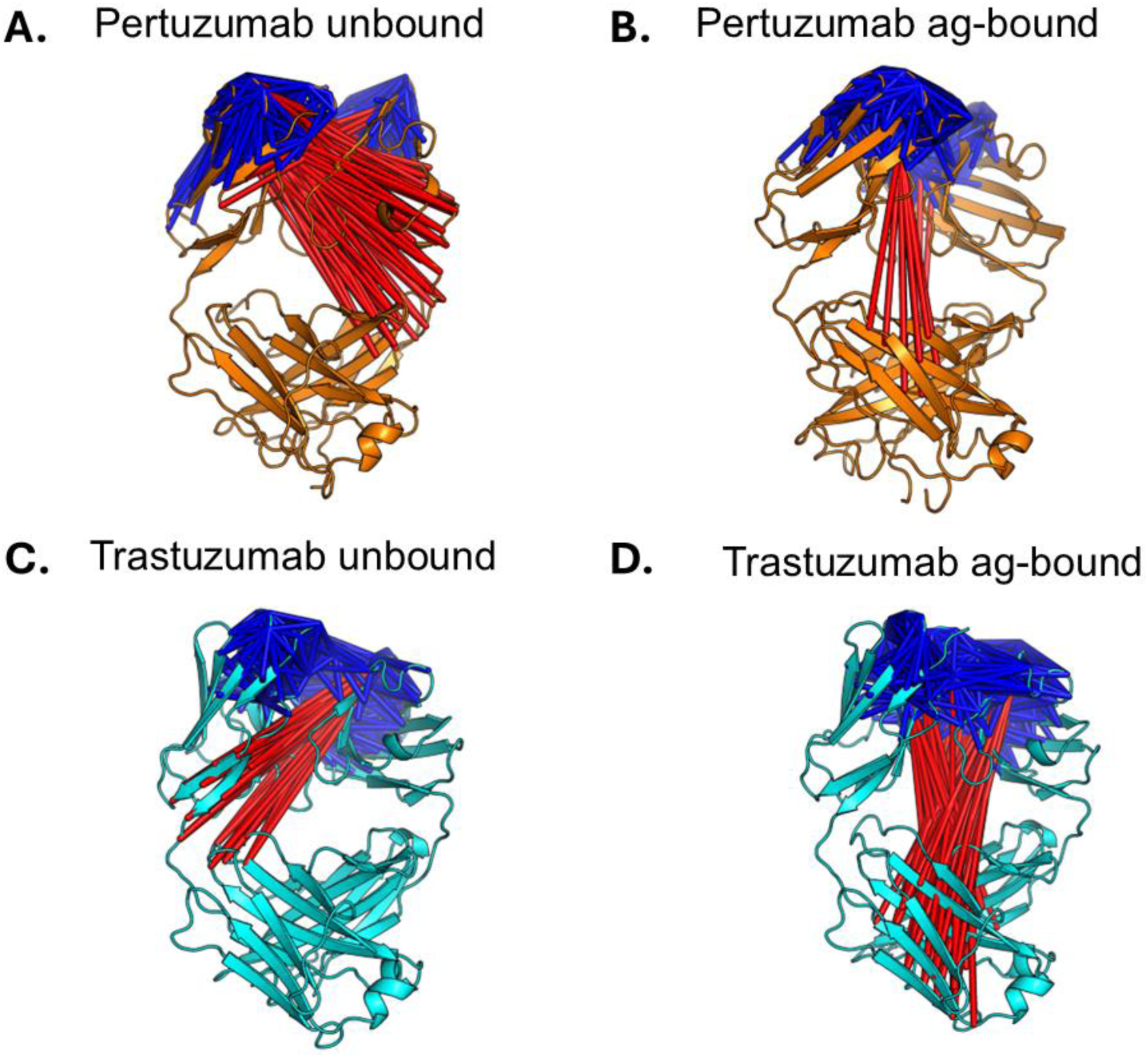
Strong cross-correlations involving CDR residues. The positive (blue) and negative (red) correlations with intensity over 0.8 involving CDR residues are shown for Pertuzumab under unbound (A) and antigen-bound (B) simulations and for Trastuzumab under unbound (C) and antigen-bound (D) simulations.

## Discussion

CDRs are the main target of optimization strategies during antibody design due to their role in antigen recognition.^36^ However, growing evidence has shown the importance of considering the framework for antibody engineering and that modifications such as CDR grafting and single-point mutations can impact the antibody’s stability.^18,19^ We hypothesize that this impact is related to perturbations in the intrinsic CDR-FW residue cross-correlation network, which affects protein dynamics and could disturb antigen binding affinities, structural stability, or other features that are important to antibody functionality. To address this, we focused on Pertuzumab and Trastuzumab, two therapeutic antibodies with identical frameworks and constant regions but distinct paratopes, allowing us to directly investigate the influence of CDR variability. Our findings are one of the first evidence of global effects of the paratope on Fab dynamics and provide new conformational insights into antibody humanization that go beyond the traditional sequence-based homology strategy.

The consistent difference in domain orientation observed in RMSD analyses was the first evidence of CDR influence in conformational dynamics that enabled further investigation of their structural impact. The angles we calculated aimed to describe the type of interdomain movement, which revealed differences between antibodies. The EH angle was consistently different between Pertuzumab and Trastuzumab in antigen-bound and unbound conditions. As this elbow is adjacent to the CDRH3, the EH angle may suffer direct structural and dynamic effects from this loop. CDRH3 is the most variable loop in antibodies because it includes an additional gene (*Diversity* - *D*) during the recombination process to form it. Thus, it contains the core residues to distinguish paratopes.^37,38^ So, it is reasonable to infer that the EH angle is the most impacted by CDR variability.

Indeed, several studies have highlighted the importance of calculating variable domain orientation for antibody design, often using ABangle.^20,21,39,40^ Because of this, we compared the angles involving constant domains to those from the reference method, and we observed that angles that account for constant domains show more statistical differences between Pertuzumab and Trastuzumab. Taken together, these findings support the recommendation of complementing the analyses of antibodies’ variable domains orientation with angles that consider constant domains in order to fully characterize the antibodies’ dynamics.

Our analysis of inter-chain contacts provides further evidence of the CDRs’ influence on antibody dynamics. MM/PBSA calculations revealed that only Pertuzumab hasCDR residues that contribute significantly to interchain stability, corroborating with previous evidence.^41,42^ Notably, Trastuzumab displayed TRP 110 (FW4) as a significant contributor to inter-chain interactions, a feature not observed in Pertuzumab. As these antibodies differ only in their CDRs, this suggests that specific paratopes may propagate distinct structural impacts to conserved framework residues, favoring or disfavoring certain FW-FW contacts. This, in turn, may lead Trastuzumab to a more restrict interdomain mobility, as it contains extra FW-FW interactions observed in both antigen-bound and unbound simulations. Regarding the antigen presence, Pertuzumab was more impacted by it, decreasing the number of CDR residues that contribute to inter-chain interactions, as calculated by MM/PBSA.

Considering all structural effects driven by different CDRs, we directly investigated the internal CDR-FW network through cross-correlation analysis. Surprisingly, the cross-correlation profile of Pertuzumab and Trastuzumab involving elbow residues in the unbound condition showed the opposite behavior from each other. In the antigen-bound condition, Pertuzumab displayed strong correlations between its CDRs and the constant domain of the same chain, whereas Trastuzumab showed strong correlations with the constant domain of the opposite chain. These findings align with the hypothesis of specific CDR-FW networks and further indicate they react differently to antigen binding. Such modulation may also have functional implications for Fc receptor interactions, as previously reported in studies that highlighted the role of Fab portion on such binding.^43–45^ More specifically, these different communication networks must alter the allosteric pathways, which could elucidate experimental differences in Fc receptor binding affinities of Pertuzumab and Trastuzumab.^18^ So, it underscores the importance of carefully evaluating the CDR-FW network during antibody design.

It is important to emphasize that the framework used in this case study successfully supports both CDR combinations of Pertuzumab and Trastuzumab, as these antibodies are already approved for clinical use. However, cases in the literature, such as non-functional or non-expressed CDR-FW combinations, highlight how perturbations in CDR-FW networks can disrupt antibody production and functionality.^17,18^ Therefore, understanding these networks is essential for ensuring compatibility between frameworks and CDRs during the antibody design process, as imbalances may compromise the delicate interplay required for functional monoclonal antibodies.

## CONCLUSION

In this study, we found that CDRs can influence several structural dynamics parameters and internal communication networks of the whole Fab of Pertuzumab and Trastuzumab. As both antibodies have the same framework and constant domains, we concluded that CDRs impact the interdomain mobility, orientation (mainly the elbow angle of the heavy chain), and interchain interactions, as observed across all MD simulations in both antigen-bound and unbound conditions. This influence is due to the CDR-FW correlation network, directly observed here, which specifically modulates each antibody’s dynamics and represents one of the first evidence of the interplay between the paratope and conserved regions. These findings emphasize the importance of performing EH angle and residue cross correlation analysis to better characterize the effects of CDRs over Fab dynamics during the antibody optimization process.

## FUNDING DETAILS

This study was financed by the São Paulo Research Foundation (FAPESP), Brasil, under Grant Process Number #2024/1665-4, the Coordenação de Aperfeiçoamento de Pessoal de Nível Superior - Brasil (CAPES) – Finance Code 001, and the Conselho Nacional de Desenvolvimento Científico e Tecnológico (CNpq), Brasil, under Grant Process Number #174494/2024-6.

## DISCLOSURE STATEMENT

The authors report that there are no competing interests to declare.

## Notes

### Competing Interest Statement

The authors have declared no competing interest.

