## Supplementary data for "Are frameworks independent from CDRs in antibodies? Exploring CDR-framework correlation networks of antibodies"

**Supplementary Table S1.** Metrics of RMSD distribution of intradomain and interdomain calculations in Pertuzumab and Trastuzumab MD simulations under antigen1bound and unbound conditions.

| | | | Pertuzumab | | | | | Trastuzumab | | | | |
|---|---|---|---|---|---|---|---|---|---|---|---|---|
| **Condition** | **Calculation** | **Domains*** | **Median** | **Mean** | **Std** | **Min** | **Max** | **Median** | **Mean** | **Std** | **Min** | **Max** |
| Antigen1bound | Intradomain | CL1CL | 1.08 | 1.10 | 0.21 | 0.62 | 2.17 | 1.04 | 1.06 | 0.18 | 0.62 | 1.74 |
| | | VL1VL | 0.52 | 0.53 | 0.07 | 0.33 | 0.99 | 0.54 | 0.55 | 0.08 | 0.34 | 0.87 |
| | | CH1CH | 0.95 | 1.01 | 0.28 | 0.50 | 2.28 | 0.84 | 0.86 | 0.18 | 0.47 | 1.98 |
| | | VH1VH | 0.61 | 0.62 | 0.10 | 0.38 | 1.14 | 0.55 | 0.56 | 0.08 | 0.32 | 1.02 |
| | Interdomain | CL1VL | 2.69 | 3.22 | 1.86 | 0.61 | 12.41 | 1.97 | 2.20 | 1.03 | 0.52 | 8.52 |
| | | VL1CL | 2.94 | 3.48 | 1.92 | 0.86 | 14.35 | 2.39 | 2.78 | 1.43 | 0.83 | 10.28 |
| | | CH1VH | 2.60 | 3.22 | 1.99 | 0.62 | 14.34 | 2.06 | 2.38 | 1.24 | 0.52 | 9.82 |
| | | VH1CH | 2.56 | 3.10 | 1.74 | 0.74 | 14.08 | 2.05 | 2.43 | 1.30 | 0.76 | 9.27 |
| Unbound | Intradomain | CL1CL | 0.98 | 1.00 | 0.22 | 0.52 | 1.72 | 0.91 | 0.97 | 0.26 | 0.49 | 2.33 |
| | | VL1VL | 0.56 | 0.59 | 0.12 | 0.34 | 1.37 | 0.54 | 0.55 | 0.08 | 0.34 | 0.94 |
| | | CH1CH | 0.99 | 1.03 | 0.23 | 0.53 | 1.77 | 0.94 | 1.00 | 0.27 | 0.49 | 2.42 |
| | | VH1VH | 0.74 | 0.79 | 0.22 | 0.39 | 1.85 | 0.69 | 0.77 | 0.26 | 0.38 | 2.30 |
| | Interdomain | CL1VL | 2.84 | 3.23 | 1.69 | 0.67 | 11.68 | 2.18 | 2.47 | 1.29 | 0.47 | 8.72 |
| | | VL1CL | 3.52 | 3.98 | 1.98 | 0.81 | 13.13 | 2.43 | 2.98 | 1.74 | 0.74 | 11.82 |
| | | CH1VH | 2.79 | 3.15 | 1.62 | 0.69 | 11.22 | 2.10 | 2.36 | 1.20 | 0.49 | 8.30 |
| | | VH1CH | 2.81 | 3.14 | 1.44 | 0.80 | 10.02 | 1.69 | 1.83 | 0.71 | 0.66 | 5.39 |

*: The prefix is the domain of fit and the suffix is the domain that was calculated.

**Supplementary Table S2.** Statistical comparison of angles involving constant domains in all simulations.

| Angle | | System | | | |
|---|---|---|---|---|---|
| | | Per unbound | Per ag-bound | Traas unbound | Traas ag-bound |
| VL-CH | Per unbound | 1 | 0.35 * | 0.19 * | NA |
| | Per ag-bound | | 1 | NA | 0.47 * |
| | Traas unbound | | | 1 | 0.10 * |
| | Traas ag-bound | | | | 1 |
| | | Per unbound | Per ag-bound | Traas unbound | Traas ag-bound |
| VH-CL | Per unbound | 1 | 0.37 * | 0.15 * | NA |
| | Per ag-bound | | 1 | NA | 0.39 * |
| | Traas unbound | | | 1 | 0.16 * |
| | Traas ag-bound | | | | 1 |
| | | Per unbound | Per ag-bound | Traas unbound | Traas ag-bound |
| EL | Per unbound | 1 | 0.32 * | 0.28 * | NA |
| | Per ag-bound | | 1 | NA | 0.22 * |
| | Traas unbound | | | 1 | 0.07 * |
| | Traas ag-bound | | | | 1 |
| | | Per unbound | Per ag-bound | Traas unbound | Traas ag-bound |
| EH | Per unbound | 1 | 0.09 * | 0.37 * | NA |
| | Per ag-bound | | 1 | NA | 0.39 * |
| | Traas unbound | | | 1 | 0.05 * |
| | Traas ag-bound | | | | 1 |

*: p-value < 0.0001; NA: Non-applicable comparison; All values were calculated according to the Kolmogorov-Smirnov test.

**Supplementary Table S3.** Spearman correlations between all angles analyzed for Pertuzumab.

| Condition | | Correlations | | | | | | | | | |
|---|---|---|---|---|---|---|---|---|---|---|---|
| | | VLCH | VHCL | EL | EH | HL | HC1 | HC2 | LC1 | LC2 | DC |
| | VLCH | 1 | -0.91 | - | -0.71 | - | - | - | - | - | - |
| | VHCL | -0.91 | 1 | -0.42 | 0.71 | - | - | - | - | - | - |
| | EL | - | -0.42 | 1 | - | - | - | - | - | - | - |
| | EH | -0.71 | 0.71 | - | 1 | - | - | - | - | - | - |
| Unbound | HL | - | - | - | - | 1 | - | - | - | - | - |
| | HC1 | - | - | - | - | - | 1 | - | - | - | -0.45 |
| | HC2 | - | - | - | - | - | - | 1 | - | - | - |
| | LC1 | - | - | - | - | - | - | - | 1 | - | - |
| | LC2 | - | - | - | - | - | - | - | - | 1 | 0.5 |
| | DC | - | - | - | - | - | -0.45 | - | - | 0.5 | 1 |
| | | VL-CH | VH-CL | EL | EH | HL | HC1 | HC2 | LC1 | LC2 | DC |
| | VLCH | 1 | -0.84 | - | -0.66 | - | - | - | - | - | - |
| | VHCL | -0.84 | 1 | - | 0.79 | - | - | - | - | - | - |
| | EL | - | - | 1 | - | - | - | - | - | - | - |
| | EH | -0.66 | 0.79 | - | 1 | - | - | - | - | - | - |
| Antigen1bound | HL | - | - | - | - | 1 | - | - | 0.63 | 0.55 | - |
| | HC1 | - | - | - | - | - | 1 | - | - | - | -0.78 |
| | HC2 | - | - | - | - | - | - | 1 | - | - | - |
| | LC1 | - | - | - | - | 0.63 | - | - | 1 | - | -0.48 |
| | LC2 | - | - | - | - | 0.55 | - | - | - | 1 | - |
| | DC | - | - | - | - | - | -0.78 | - | -0.48 | - | 1 |

Interdomain angles: VLCH, VHCL, EL and EH; ABangle: HL, HC1, HC2, LC1, LC2 and DC.

**Supplementary Table S4.** Spearman correlations between all angles analyzed for Trastuzumab.

| Condition | | Correlations | | | | | | | | | |
|---|---|---|---|---|---|---|---|---|---|---|---|
| | | VL-CH | VH-CL | EL | EH | HL | HC1 | HC2 | LC1 | LC2 | DC |
| $HER2^-$ | VL-CH | 1 | -0.9 | - | -0.74 | - | - | - | - | - | - |
| | VH-CL | -0.9 | 1 | -0.42 | 0.69 | - | - | - | - | - | - |
| | EL | - | -0.42 | 1 | - | - | - | - | - | - | - |
| | EH | -0.74 | 0.69 | - | 1 | - | - | - | - | - | - |
| | HL | - | - | - | - | 1 | - | - | - | 0.6 | - |
| | HC1 | - | - | - | - | - | 1 | - | - | - | - |
| | HC2 | - | - | - | - | - | - | 1 | - | - | - |
| | LC1 | - | - | - | - | - | - | - | 1 | - | - |
| | LC2 | - | - | - | - | 0.6 | - | - | - | 1 | - |
| | DC | - | - | - | - | - | - | - | - | - | 1 |
| | | VL-CH | VH-CL | EL | EH | HL | HC1 | HC2 | LC1 | LC2 | DC |
| $Her2^+$ | VL-CH | 1 | -0.81 | - | -0.6 | - | - | - | - | - | - |
| | VH-CL | -0.81 | 1 | - | 0.64 | - | - | - | - | - | - |
| | EL | - | - | 1 | - | - | - | - | - | - | - |
| | EH | -0.6 | 0.64 | - | 1 | - | - | - | - | - | - |
| | HL | - | - | - | - | 1 | - | - | - | - | - |
| | HC1 | - | - | - | - | - | 1 | 0.65 | - | - | - |
| | HC2 | - | - | - | - | - | 0.65 | 1 | - | - | - |
| | LC1 | - | - | - | - | - | - | - | 1 | - | - |
| | LC2 | - | - | - | - | - | - | - | - | 1 | - |
| | DC | - | - | - | - | - | - | - | - | - | 1 |

Interdomain angles: VLCH, VHCL, EL and EH; ABangle: HL, HC1, HC2, LC1, LC2 and DC.

**Supplementary Table S5.** Statistical comparison of angles calculated by ABangle in all simulations.

| Angle | | System | | | |
|---|---|---|---|---|---|
| | | Per unbound | Per ag-bound | Traas unbound | Traas ag-bound |
| HL | Per unbound | 1 | 0.56 * | 0.14 * | NA |
| | Per ag-bound | | 1 | NA | 0.13 * |
| | Traas unbound | | | 1 | 0.50 * |
| | Traas ag-bound | | | | 1 |
| | | Per unbound | Per ag-bound | Traas unbound | Traas ag-bound |
| HC1 | Per unbound | 1 | 0.67 * | 0.23 * | NA |
| | Per ag-bound | | 1 | NA | 0.11 * |
| | Traas unbound | | | 1 | 0.38 * |
| | Traas ag-bound | | | | 1 |
| | | Per unbound | Per ag-bound | Traas unbound | Traas ag-bound |
| HC2 | Per unbound | 1 | 0.59 * | 0.30 * | NA |
| | Per ag-bound | | 1 | NA | 0.33 * |
| | Traas unbound | | | 1 | 0.51 * |
| | Traas ag-bound | | | | 1 |
| | | Per unbound | Per ag-bound | Traas unbound | Traas ag-bound |
| LC1 | Per unbound | 1 | 0.30 * | 0.20 * | NA |
| | Per ag-bound | | 1 | NA | 0.13 * |
| | Traas unbound | | | 1 | 0.50 * |
| | Traas ag-bound | | | | 1 |
| | | Per unbound | Per ag-bound | Traas unbound | Traas ag-bound |
| LC2 | Per unbound | 1 | 0.64 * | 0.21 * | NA |
| | Per ag-bound | | 1 | NA | 0.20 * |
| | Traas unbound | | | 1 | 0.71 * |
| | Traas ag-bound | | | | 1 |
| | | Per unbound | Per ag-bound | Traas unbound | Traas ag-bound |
| DC | Per unbound | 1 | 0.41 * | 0.18 * | NA |
| | Per ag-bound | | 1 | NA | 0.30 * |
| | Traas unbound | | | 1 | 0.47 * |
| | Traas ag-bound | | | | 1 |

*: p-value < 0.0001; NA: Non-applicable comparison; All values were calculated according to the Kolmogorov-Smirnov test.

**Supplementary Table S6.** Contacts of MM/PBSA relevant residues of Pertuzumab simulations in unbound condition.

| Residue* | VDW | EEL | EPB | Total ΔG | Ab segment | Contacts# |
|---|---|---|---|---|---|---|
| K TYR 91 | -3.65 | 0.29 | 0.28 | -3.09 | cdr3 | PHE 99A - cdr3 (98.62%), TYR 99B - cdr3 (71.81%), SER 99 - cdr3 (53.17%), PRO 98 - cdr3 (0.58%) |
| K PHE 98 | -4.63 | -0.83 | 1.41 | -4.05 | fw4 | LEU 45 - fw2 (99.02%), PHE 100 - cdr3 (76.58%), TRP 47 - fw2 (64.21%), GLU 46 - fw2 (51.55%), TRP 103 - fw4 (46.65%), VAL 37 - fw2 (43.51%), PHE 99A - cdr3 (10.08%), GLY 44 - fw2 (0.10%) |
| K PHE 118 | -5.18 | -0.29 | 1.07 | -4.4 | constant | LEU 124 - constant (94.10%), ALA 125 - constant (79.82%), ALA 137 - constant (74.09%), PRO 126 - constant (63.23%), LEU 138 - constant (43.73%), VAL 181 - constant (38.07%), GLY 139 - constant (19.36%), SER 127 - constant (4.56%), SER 130 - constant (0.62%), LEU 141 - constant (0.04%), ALA 136 - constant (0.02%) |
| H GLN 39 | -0.93 | -7.56 | 5.63 | -2.86 | fw2 | GLN 38 - fw2 (98.30%), TYR 87 - fw3 (81.50%), PRO 44 - fw2 (3.54%), LYS 103 - fw4 (0.02%) |
| H LEU 45 | -5.25 | -0.35 | 2.21 | -3.4 | fw2 | PHE 98 - fw4 (99.02%), PRO 44 - fw2 (93.34%), TYR 87 - fw3 (66.77%), GLN 3 - fw1 (9.64%), GLN 38 - fw2 (7.30%), GLY 99 - fw4 (2.70%), TYR 36 - fw2 (0.48%), GLN 100 - fw4 (0.04%) |
| H TRP 47 | -3.42 | 0.14 | 0.44 | -2.84 | fw2 | TYR 94 - cdr3 (81.22%), TYR 96 - cdr3 (72.37%), PHE 98 - fw4 (64.21%), PRO 95 - cdr3 (54.31%), GLN 89 - cdr3 (0.30%) |
| H PHE 99[a] | -4.67 | -0.99 | 3.1 | -2.56 | cdr3 | TYR 91 - cdr3 (98.62%), TYR 96 - cdr3 (71.23%), GLN 89 - cdr3 (66.47%), TYR 49 - fw2 (19.46%), PHE 98 - fw4 (10.08%), LEU 46 - fw2 (5.24%), ALA 34 - fw2 (1.22%), TYR 94 - cdr3 (0.92%), TYR 36 - fw2 (0.76%), SER 50 - cdr2 (0.44%), GLN 90 - cdr3 (0.40%), TYR 92 - cdr3 (0.02%) |
| H TYR 99B | -5.96 | 0.6 | 1.5 | -3.87 | cdr3 | LEU 46 - fw2 (97.40%), TYR 36 - fw2 (73.27%), TYR 49 - fw2 (71.91%), TYR 91 - cdr3 (71.81%), GLN 89 - cdr3 (38.27%), ALA 34 - fw2 (35.41%), TYR 55 - fw3 (25.67%), THR 56 - fw3 (14.08%), SER 50 - cdr2 (0.46%) |
| H LEU 130 | -3.17 | -0.72 | 1.24 | -2.66 | constant | PHE 118 - constant (94.10%), VAL 133 - constant (84.54%), SER 121 - constant (13.02%), LEU 135 - constant (9.04%), PRO 119 - constant (4.70%), GLN 124 - constant (2.68%), SER 131 - constant (0.10%), PRO 120 - constant (0.04%) |
| H PHE 172 | -6.09 | -0.55 | 1.73 | -4.91 | constant | SER 176 - constant (96.70%), SER 174 - constant (88.38%), LEU 135 - constant (85.82%), LEU 175 - constant (84.40%), THR |

| Residue* | VDW | EEL | EPB | Total ΔG | Ab segment | Contacts[#] |
|---|---|---|---|---|---|---|
| | | | | | | 164 - constant (77.26%), SER 162 - constant (63.33%), VAL 163 - constant (59.75%), ASN 137 - constant (19.58%), LEU 136 - constant (19.26%), GLU 165 - constant (10.56%), PHE 116 - constant (7.86%) |
| H PRO 173 | -2.88 | -1.6 | 1.4 | -3.08 | constant | VAL 163 - constant (86.18%), THR 164 - constant (80.42%), SER 162 - constant (68.61%), GLU 165 - constant (34.79%), SER 176 - constant (19.92%), SER 174 - constant (17.64%), LEU 175 - constant (15.42%) |

*: Residues are described by their chain (kappa or heavy, K or H, respectively), name, and number in PDB; [#]: Contact residues are described by their name, number, antibody segment, and frequency of contact time; VDW, EEL, and EPB represent Van der Waals, electrostatic, and solvation contributions, respectively, as calculated in the MM/PBSA analysis.

**Supplementary Table S7.** Contacts of MM/PBSA relevant residues of Pertuzumab simulations in antigen-bound condition.

| Residue* | VDW | EEL | EPB | Total ΔG | Ab segment | Contacts# |
|---|---|---|---|---|---|---|
| K PHE 98 | -4.76 | -0.64 | 1.34 | -4.05 | fw4 | LEU 45 - fw2 (97.94%), TRP 47 - fw2 (80.74%), PHE 100 - cdr3 (72.07%), GLU 46 - fw2 (66.85%), VAL 37 - fw2 (46.31%), TRP 103 - fw4 (30.87%), PHE 99A - cdr3 (3.96%), GLY 44 - fw2 (0.02%) |
| K PHE 116 | -3.4 | -0.2 | 0.54 | -3.05 | constant | ALA 136 - constant (89.68%), THR 135 - constant (82.96%), ALA 137 - constant (71.49%), THR 183 - constant (31.15%), SER 132 - constant (14.78%), THR 131 - constant (8.00%), SER 130 - constant (6.36%), PRO 126 - constant (2.38%), VAL 181 - constant (0.08%), GLY 133 - constant (0.02%) |
| K PHE 118 | -5.63 | -0.36 | 1.15 | -4.85 | constant | LEU 124 - constant (94.52%), ALA 137 - constant (93.26%), ALA 125 - constant (79.04%), PRO 126 - constant (74.89%), LEU 138 - constant (61.49%), VAL 181 - constant (45.89%), GLY 139 - constant (13.06%), SER 127 - constant (0.12%), LYS 214 - constant (0.04%) |
| H LEU 45 | -5.07 | -0.05 | 1.84 | -3.28 | fw2 | PHE 98 - fw4 (97.94%), PRO 44 - fw2 (81.78%), TYR 87 - fw3 (65.65%), GLN 38 - fw2 (9.12%), GLY 99 - fw4 (6.30%), GLN 100 - fw4 (0.24%), TYR 36 - fw2 (0.06%) |
| H TRP 47 | -4.15 | 0.16 | 0.57 | -3.43 | fw2 | TYR 96 - cdr3 (94.94%), TYR 94 - cdr3 (93.62%), PHE 98 - fw4 (80.74%), PRO 95 - cdr3 (77.72%), GLN 89 - cdr3 (1.32%), THR 97 - cdr3 (0.08%) |
| H TYR 99B | -5.59 | 1.19 | 1.47 | -2.92 | cdr3 | LEU 46 - fw2 (96.52%), TYR 49 - fw2 (79.94%), TYR 91 - cdr3 (73.81%), TYR 36 - fw2 (71.81%), ALA 34 - fw2 (31.73%), GLN 89 - cdr3 (18.64%), TYR 55 - fw3 (13.48%), THR 56 - fw3 (8.18%), SER 50 - cdr2 (4.72%) |
| H PHE 128 | -2.86 | -1.11 | 1.46 | -2.51 | constant | SER 121 - constant (74.75%), GLN 124 - constant (74.31%), GLU 123 - constant (56.49%), PRO 120 - constant (7.76%), PRO 119 - constant (5.58%), SER 127 - constant (4.66%), SER 131 - constant (1.64%), THR 129 - constant (1.18%), ASP 122 - constant (0.30%), LYS 126 - constant (0.26%), VAL 133 - constant (0.14%) |
| H LEU 130 | -3.16 | -0.62 | 1.23 | -2.56 | constant | PHE 118 - constant (94.52%), VAL 133 - constant (76.30%), SER 121 - constant (16.80%), PRO 119 - constant (8.96%), GLN 124 - constant (3.32%), LEU 135 - constant (1.32%), PRO 120 - constant (1.00%), SER 131 - constant (0.36%) |

| Residue* | VDW | EEL | EPB | Total ΔG | Ab segment | Contacts# |
|---|---|---|---|---|---|---|
| H PHE 172 | -6.44 | -0.77 | 2 | -5.21 | constant | THR 164 - constant (97.38%), LEU 175 - constant (97.08%), SER 176 - constant (95.08%), SER 162 - constant (92.00%), VAL 163 - constant (90.96%), SER 174 - constant (85.80%), LEU 135 - constant (77.48%), LEU 136 - constant (1.00%) |

*: Residues are described by their chain (kappa or heavy, K or H, respectively), name, and number in PDB; #: Contact residues are described by their name, number, antibody segment, and frequency of contact time; VDW, EEL, and EPB represent Van der Waals, electrostatic, and solvation contributions, respectively, as calculated in the MM/PBSA analysis.

**Supplementary Table S8.** Contacts of MM/PBSA relevant residues of Trastuzumab simulations in unbound condition.

| Residue* | VDW | EEL | EPB | Total ΔG | Ab segment | Contacts# |
|---|---|---|---|---|---|---|
| K GLN 38 | -0.97 | -7.42 | 5.86 | -2.54 | fw2 | GLN 39 - fw2 (97.62%), TYR 95 - fw3 (80.42%), LEU 45 - fw2 (9.96%) |
| K PHE 98 | -4.28 | -0.23 | 0.87 | -3.64 | fw4 | LEU 45 - fw2 (96.42%), TRP 110 - fw4 (77.78%), MET 107 - cdr3 (72.91%), TRP 47 - fw2 (57.53%), GLU 46 - fw2 (38.97%), VAL 37 - fw2 (32.51%) |
| K PHE 116 | -3.07 | -0.46 | 0.8 | -2.73 | constant | ALA 144 - constant (79.18%), ALA 143 - constant (61.57%), THR 142 - constant (59.79%), THR 190 - constant (49.87%), VAL 188 - constant (13.16%), PHE 173 - constant (4.40%), SER 139 - constant (2.96%), THR 138 - constant (0.74%), PRO 133 - constant (0.30%), SER 134 - constant (0.26%), SER 135 - constant (0.06%), HIS 171 - constant (0.06%) |
| K PHE 118 | -5.72 | -1.47 | 2.23 | -4.96 | constant | LEU 131 - constant (97.12%), ALA 144 - constant (86.46%), ALA 132 - constant (80.14%), LEU 145 - constant (61.83%), PRO 133 - constant (58.05%), VAL 188 - constant (56.55%), GLY 146 - constant (22.16%), SER 134 - constant (7.96%), SER 135 - constant (2.60%), PHE 129 - constant (0.02%) |
| H LEU 45 | -4.83 | -0.35 | 2.21 | -2.97 | fw2 | PHE 98 - fw4 (96.42%), PRO 44 - fw2 (86.96%), TYR 87 - fw3 (53.47%), GLN 38 - fw2 (9.96%), GLY 99 - fw4 (0.10%) |
| H TRP 47 | -3.73 | -0.55 | 1.06 | -3.22 | fw2 | PRO 95 - cdr3 (99.52%), PRO 96 - cdr3 (99.28%), THR 94 - cdr3 (63.65%), PHE 98 - fw4 (57.53%) |
| H TRP 110 | -5.33 | -0.13 | 1.44 | -4.03 | fw4 | TYR 36 - fw2 (99.24%), PRO 44 - fw2 (88.94%), PHE 98 - fw4 (77.78%), ALA 43 - fw2 (58.81%), LYS 45 - fw2 (15.80%), LEU 46 - fw2 (3.58%) |
| H PHE 129 | -3.29 | -1.02 | 1.4 | -2.91 | constant | GLN 124 - constant (85.72%), GLU 123 - constant (69.15%), SER 121 - constant (64.51%), SER 127 - constant (19.36%), SER 131 - constant (8.22%), PRO 120 - constant (3.20%), ALA 130 - constant (2.78%), PRO 119 - constant (2.30%), CYS 214 - constant (0.30%), PHE 118 - constant (0.02%), ASP 122 - constant (0.02%), LYS 126 - constant (0.02%), THR 129 - constant (0.02%), VAL 132 - constant (0.02%), VAL 133 - constant (0.02%) |
| H LEU 131 | -3.54 | -0.5 | 1.43 | -2.61 | constant | PHE 118 - constant (97.12%), VAL 133 - constant (91.98%), SER 121 - constant (12.44%), GLN 124 - constant (8.28%), LEU 135 - constant (7.86%), PRO 119 - constant (4.98%), SER 131 - constant (1.20%), PRO 120 - constant (0.42%), VAL 132 - constant (0.18%) |

| Residue* | VDW | EEL | EPB | Total ΔG | Ab segment | Contacts# |
|---|---|---|---|---|---|---|
| H PHE 173 | -6 | -1 | 1.97 | -5.03 | constant | SER 176 - constant (97.98%), SER 174 - constant (93.20%), LEU 175 - constant (89.30%), LEU 135 - constant (87.50%), THR 164 - constant (65.19%), VAL 163 - constant (56.69%), SER 162 - constant (55.75%), LEU 136 - constant (34.03%), ASN 137 - constant (26.47%), PHE 116 - constant (4.40%) |
| H PRO 174 | -2.65 | -0.73 | 0.87 | -2.51 | constant | THR 164 - constant (85.24%), VAL 163 - constant (83.60%), SER 162 - constant (74.95%), SER 176 - constant (20.08%), SER 174 - constant (9.66%), LEU 175 - constant (9.32%), GLU 165 - constant (3.90%) |

*: Residues are described by their chain (kappa or heavy, K or H, respectively), name, and number in PDB; #: Contact residues are described by their name, number, antibody segment, and frequency of contact time; VDW, EEL, and EPB represent Van der Waals, electrostatic, and solvation contributions, respectively, as calculated in the MM/PBSA analysis.

**Supplementary Table S9.** Contacts of MM/PBSA relevant residues of Trastuzumab simulations in antigen-bound condition.

| Residue* | VDW | EEL | EPB | Total ΔG | Ab segment | Contacts# |
|---|---|---|---|---|---|---|
| K GLN 38 | -0.83 | -8.55 | 6.41 | -2.98 | fw2 | GLN 39 - fw2 (99.58%), TYR 95 - fw3 (82.48%), LEU 45 - fw2 (3.78%) |
| K PHE 98 | -4.26 | -0.25 | 0.9 | -3.61 | fw4 | LEU 45 - fw2 (96.60%), TRP 47 - fw2 (72.13%), TRP 110 - fw4 (62.33%), MET 107 - cdr3 (53.75%), GLU 46 - fw2 (48.79%), VAL 37 - fw2 (38.13%) |
| K PHE 116 | -2.88 | -0.57 | 0.79 | -2.66 | constant | ALA 143 - constant (72.07%), ALA 144 - constant (67.89%), THR 142 - constant (43.67%), PRO 133 - constant (14.86%), THR 190 - constant (14.22%), SER 137 - constant (7.84%), THR 138 - constant (4.02%), VAL 188 - constant (3.00%), SER 134 - constant (1.96%), SER 139 - constant (0.30%), LEU 196 - constant (0.04%), HIS 171 - constant (0.02%) |
| K PHE 118 | -5.45 | -1.37 | 2.15 | -4.67 | constant | LEU 131 - constant (96.18%), ALA 144 - constant (86.24%), ALA 132 - constant (79.62%), PRO 133 - constant (72.05%), LEU 145 - constant (62.59%), VAL 188 - constant (35.17%), GLY 146 - constant (9.00%), LYS 136 - constant (0.04%), PHE 129 - constant (0.02%), SER 134 - constant (0.02%) |
| H LEU 45 | -4.91 | -0.48 | 2.34 | -3.05 | fw2 | PHE 98 - fw4 (96.60%), PRO 44 - fw2 (84.06%), TYR 87 - fw3 (56.73%), GLN 3 - fw1 (11.58%), GLN 38 - fw2 (3.78%), GLY 99 - fw4 (0.20%) |
| H TRP 47 | -3.93 | -0.54 | 1.29 | -3.17 | fw2 | PRO 95 - cdr3 (98.88%), PRO 96 - cdr3 (94.46%), PHE 98 - fw4 (72.13%), THR 94 - cdr3 (59.77%) |
| H TRP 110 | -5.33 | 0.58 | 1.47 | -3.28 | fw4 | TYR 36 - fw2 (97.54%), PRO 44 - fw2 (88.06%), PHE 98 - fw4 (62.33%), ALA 43 - fw2 (59.45%), LYS 45 - fw2 (30.31%), LEU 46 - fw2 (3.54%) |
| H PHE 129 | -3.34 | -0.82 | 1.39 | -2.77 | constant | GLN 124 - constant (84.16%), SER 121 - constant (79.16%), GLU 123 - constant (66.35%), SER 127 - constant (9.58%), PRO 120 - constant (8.84%), SER 131 - constant (8.76%), ALA 130 - constant (8.40%), PRO 119 - constant (1.04%), VAL 133 - constant (0.38%), PHE 118 - constant (0.02%), ASP 122 - constant (0.02%) |
| H LEU 131 | -3.5 | -0.58 | 1.43 | -2.66 | constant | PHE 118 - constant (96.18%), VAL 133 - constant (95.76%), SER 121 - constant (6.74%), LEU 135 - constant (6.48%), PRO 119 - constant (4.80%), GLN 124 - constant (2.60%), PRO 120 - constant (0.28%), SER 131 - constant (0.08%) |
| H PHE 173 | -6.1 | -0.96 | 1.97 | -5.09 | constant | LEU 175 - constant (97.82%), SER 176 - constant (95.46%), THR 164 - constant |

| Residue* | VDW | EEL | EPB | Total ΔG | Ab segment | Contacts# |
|---|---|---|---|---|---|---|
| | | | | | | (94.72%), LEU 135 - constant (90.70%), SER 174 - constant (88.84%), VAL 163 - constant (73.15%), SER 162 - constant (72.93%), LEU 136 - constant (6.82%), ASN 137 - constant (3.30%) |

*: Residues are described by their chain (kappa or heavy, K or H, respectively), name, and number in PDB; #: Contact residues are described by their name, number, antibody segment, and frequency of contact time; VDW, EEL, and EPB represent Van der Waals, electrostatic, and solvation contributions, respectively, as calculated in the MM/PBSA analysis.
